# Do you see what I see? Diffusion imaging protocol heterogeneity biases ischemic core volume, location, and clinical associations in acute stroke

**DOI:** 10.1101/2024.03.19.585494

**Authors:** Jonathan Rafael-Patiño, Elda Fischi-Gomez, Antoine Madrona, Veronica Ravano, Bénédicte Maréchal, Tobias Kober, Silvia Pistocchi, Alexander Salerno, Guillaume Saliou, Patrik Michel, Roland Wiest, Richard McKinley, Jonas Richiardi

## Abstract

**Purpose:** Diffusion-weighted magnetic resonance imaging (DWI) is essential for diagnosing ischemic stroke and identifying targets for emergency revascularization. Apparent diffusion coefficient (ADC) maps derived from DWI are commonly used to locate the infarct core, but they are not strictly quantitative and can vary across platforms and sites due to technical factors. This retrospective study was conducted to examine how differences in ADC map generation, resulting from varied protocols across platforms and sites, affect the determination of infarct core size, location, and related clinical outcomes in acute stroke.

**Methods:** In this retrospective study, 726 acute anterior circulation stroke patients admitted to the Lausanne University Hospital between May 2018 and January 2021 were selected. DWI data were used to generate ADC maps as they would appear from different protocols: two simulated with low and medium angular resolution (4 and 12 diffusion gradient directions) and one with high angular resolution (20 directions). Using a DEFUSE like criteria and image post-processing, ischemic cores were localized; core volume, location, and associations to National Institutes of Health Stroke Scale (NIHSS) and modified Rankin Scale (mRS) scores were compared between the two imaging sequences.

**Results:** Significant differences were observed in the ADC distribution within white matter, particularly in the kurtosis and skewness, with the segmented infarct core volume being higher in protocols with reduced angular resolution compared to the 20-directions data (7.63 ml vs. 3.78 ml). The volumetric differences persisted after correcting for age, sex, and type of intervention. Infarcted voxels locations varied significantly between the two protocols. This variability affected associations between infarct core volume and clinical scores, with lower associations observed for 4-directions data compared to 20-directions data for NIHSS at admission and after 24 hours, and mRS after 3 months, further confirmed by multivariate regression.

**Conclusions:** Imaging protocol heterogeneity leads to significant changes in the ADC distribution, ischemic core location, size, and association with clinical scores. Work is needed in standardizing imaging protocols to improve the reliability of ADC as an imaging biomarker in stroke management.protocols to improve the reliability of ADC as an imaging biomarker in stroke management.

**Key Results:**

- Orientation changes in diffusion imaging significantly impact ADC distribution and threshold-based infarct core volume determination, affecting multi-centric studies.
- Lower number of directions in DWI acquisitions weakens associations between infarct volume measurements and clinical scores.
- Findings emphasize the impact of DWI acquisition protocol heterogeneity and image processing on acute stroke workups.

## Introduction

In acute stroke imaging centers, diffusion-weighted imaging (DWI) is indispensable for early and accurate diagnosis. This imaging technique facilitates the identification of emergency revascularization treatment targets, and estimates the final infarct volume, enhancing long-term patient outcomes. Current guidelines (Powers et al., 2018) advocate for the use of DWI and perfusion-weighted imaging (PWI), to diagnose acute ischemic stroke for patients with a stroke of 6-24 hours duration (known onset time) (Berge et al., 2021). DWI is effective for distinguishing between lesions of varying diffusion rates within minutes of stroke onset. In addition, it provides an estimate of the final infarct volume. These estimates form the basis for computing volumetric differences between PWI- and DWI-based segmentations, crucial for identifying potentially salvageable tissue (Rao et al., 2019). The speedy and precise delineation of this tissue is paramount in stroke management as the extent of the ischemic penumbra— the severely hypoperfused neurophysiologically silent brain tissue with persistent cellular integrity — is time sensitive.

Apparent Diffusion Coefficient (ADC) estimates hold high clinical significance in stroke management, but their reproducibility across various imaging platforms remains contentious (Sasaki et al., 2008). Variations in acquisition parameters and other influencing factors contribute to the difficulty of consistently and accurately measuring ADC in stroke scenarios (Donati et al., 2014; Galinovic et al., 2011; Habegger et al., 2018; Pistocchi et al., 2022). Despite these challenges, ADC maps are integral in clinical practice and trials for quantifying stroke lesions and have been widely investigated as prognostic biomarkers.

The present study aims to address the critical issue of ADC variability in acute stroke lesion segmentation due to protocol heterogeneity, as might be encountered between different hospitals. The study focused on the relationship between the volume of segmented lesions and the number and calibration of the diffusion encoding gradients – a prominent pitfall that affects the reproducibility of multi-center studies (Fedeli et al., 2018). More diffusion encoding gradients lead to a higher signal-to-noise ratio, which in turn improves ADC measurements. Conversely, the lack of 3D encoding gradients in certain protocols can also introduce sensitivity to head tilting during scanning, leading to increased variability in ADC measurements.

To investigate the dependence of segmented lesion volume on the number of diffusion encoding gradients, we utilized a cohort of patients who underwent imaging with a high number of diffusion vectors. We then performed physics-based simulations of ADC maps for low-resolution protocols (akin to protocols used in many clinical centers) to determine the effect of protocol heterogeneity on ADC variation. In doing so, we demonstrate the influence of this variability on the correlation with other factors, such as the NIHSS score. Our study provides insight into these relationships, contributing to a deeper understanding of the challenges and potential solutions for reducing ADC variability in acute stroke lesion segmentation across different imaging protocols and centers.

## Materials and Methods

This retrospective study was performed in line with the principles of the Declaration of Helsinki. Approval was granted by the regional Ethics Committee, including consent waiver (CER-VD 2022-00119). This section follows STROBE reporting guidelines (von Elm et al., 2008).

### Study Sample

From the observational ASTRAL registry (Michel et al., 2010), we obtained an initial cohort of 1240 visits for patients with anterior circulation acute stroke, with ages greater than 18 years and admitted to the Lausanne University Hospital between May 2018 and January 2021. We excluded patients without paired DWI and structural imaging data (N=323), patients with poor structural imaging quality (N=83), reflected in poor quality score in the segmentation procedure explained in the following sections, and patients without a high-resolution DWI imaging necessary to compute the synthetic low resolution protocol ADC maps (N=78). The final dataset included 726 patients. The patient flow diagram in Figure 1 details the patient selection. Table 1 summarises the characteristics of the patient sample. It is noteworthy that 253 patients from this cohort have been previously incorporated into several larger multi-centric studies (Boulenoir et al., 2022; Filioglo et al., 2022; Fischer et al., 2022; Marto et al., 2023; Nguyen et al., 2023; Schwarz et al., 2023), which conducted comparative analyses of CT and MRI in terms of workflow metrics, therapies, and outcomes. However, the objectives of these prior investigations differ from the current study.

**Table 1.** Breakdown of the 726 patients in the study sample. rtPA: recombinant tissue-type plasminogen activator. EVT: endovascular therapy (thrombectomy). NIHSS: National Institutes of Health Stroke Scale. mRS: modified Rankin scale.

| Intervention Type | EVT | rtPA | Bridging | none |
| --- | --- | --- | --- | --- |
| Total | 100 | 170 | 94 | 362 |
| Age, y (mean (SD)) | 72.62 (12.04) | 73.09 (13.60) | 73.84 (14.22) | 71.08 (15.31) |
| Male (%) | 64.0 | 58.2 | 45.7 | 58.3 |
| NIHSS (admission) (median [IQR]) | 12.00 [6.75, 19.00] | 5.00 [3.00, 8.00] | 11.00 [6.00, 19.00] | 2.00 [1.00, 5.00] |
| NIHSS (24h) (median [IQR]) | 9.00 [4.00, 16.25] | 3.00 [1.00, 6.00] | 6.00 [2.75, 15.25] | 2.00 [0.00, 4.00] |
| mRS (3 months) (median [IQR]) | 3.00 [2.00, 6.00] | 2.00 [1.00, 3.00] | 3.00 [2.00, 4.00] | 2.00 [1.00, 3.00] |

**Figure 1:**
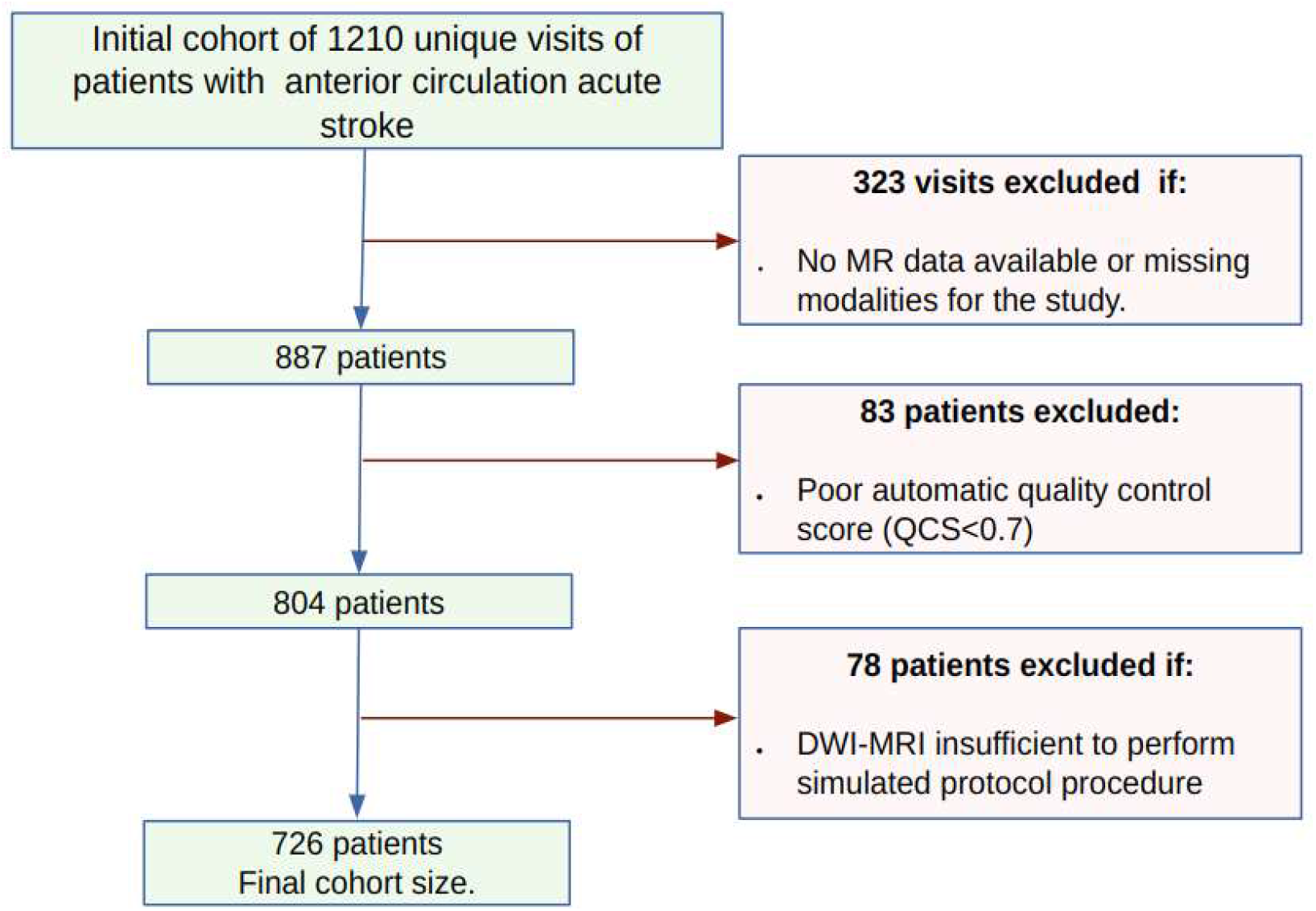
Flow chart of patient’s cohort after applying exclusion and inclusion criteria.

### Imaging protocol

Following institutional standard operating procedures, MRI was attempted as the initial stroke imaging after a standard medical history taking and a directed clinical exam by an emergency physician and a neurologist. Contraindications for MRI were respected following a prespecified checklist, and goals for door-to-imaging delay was <15 minutes for acutely treatable patients, and < 2 hours for the others.

Imaging was performed on a 3T MRI system (MAGNETOM Vida, Siemens Healthcare, Erlangen, Germany) using a 32-channels head coil. Only the DWI and structural T1- and T2-weighted images were considered in the study. The DWI acquisition was performed using a whole-brain single-shell EPI sequence with multi-band acceleration(factor 2) and included 20 diffusion encoding (b-vectors) at a maximum b-value of 1000s/mm^2^ and 10 additional *b* = 0 (*b*0) images. The structural images were acquired using a sagittal T2-weighted fluid-attenuated inversion recovery (FLAIR) turbo spin-echo (TSE) sequence. The imaging parameters were as follows: DWI (TE/TR = 80/3000 ms, resolution = 1.6×1.6×3mm^3^, a field of view (FOV) = 144×148×52 mm, flip angle (FA) = 90 degrees); structural T2-weighted (TE/TR = 87/9000 ms, inversion time (TI) = 2500 ms, resolution = 0.35×0.35×3mm^3^, FOV = 440×640×51, FA = 150 degrees).

### Image processing

The ten DWI *b*0 repetitions images per patient were averaged and used as targets for the registration with the structural images. T1 and T2 images were registered to the averaged non-weighted diffusion image 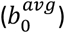 using a multi-resolution mutual information registration with b-spline interpolation (Klein et al., 2010). The total intracranial volume (TIV) was extracted from the registered T1-weighted image using Synthseg (Billot et al., 2023) a CNN-based segmentation network for brain MRI scans of any contrast and resolution without retraining. Tissue segmentation and partial volume estimates for the white matter (WM), grey matter (GM) and cerebrospinal fluid (CSF) were also obtained from Synthseg’s automatic volume estimates. The ADC maps were computed from the DWI using the mean exponential decay of the diffusion signal The ADC maps were computed from the DWI using the mean exponential decay of the diffusion signal (Mascalchi et al., 2005).

### Simulated protocol heterogeneity

Despite their widespread use for stroke workups, apparent diffusion coefficient (ADC) maps are not strictly quantitative, and these maps can be influenced by hardware and acquisition protocols (Schmeel, 2019). Given the ethical challenges of implementing repeated acquisitions with different protocols during the acute phase, we took advantage of the high angular resolution of the acquired diffusion directions to simulate variations in the acquisition. Specifically, we exploit the physics behind DWI acquisitions with high-angular resolution - that is, those employing several b-vector directions - to generate new data for each patient. This proposed technique to sub-sample the diffusion directions and create a new volume with a lower angular resolution allows us to compute a new ADC map that mimics other acquisition protocols commonly used in clinical settings. The resulting data can be used to study the impact of different DWI protocols on ADC variability and acute stroke lesion segmentation.

To improve the usefulness and realistic representation of the synthetic samples, a strategy is employed to mimic the b-vector acquisition methods of other imaging centres. This is accomplished through a greedy search that maximizes the cosine similarity between the target b-vectors and the candidate b-vectors for subsampling. The use of this search allows us to identify the candidate b-vectors that are most similar to the target b-vectors, thereby ensuring that the resulting synthetic samples accurately reflect the diffusion characteristics of different scan conditions. The cosine similarity between two vectors, U and V, is calculated as 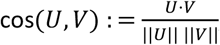 where *V* are the target vectors b-vectors and *U* are the candidate b-vectors for subsampling.

In this study, the focus was on matching the readout-segmented EPI RESOLVE diffusion protocol (Julien Cohen-Adad, 2012; Robson et al., 1997) with four b-vectors acquisition, which is commonly used in Emergency Room (ER) settings and compare it with a second sub-sampled protocol with a medium angular resolution (12 diffusion gradients) and the original high-angular resolution protocol with 20 directions. To simulate these protocols, the DWI images (originally acquired with 20 directions) were sub-sampled into non-overlapping sets of three orthogonal directions plus one additional direction (4-directions scheme, Figure 2 panel A), and 12 equispaced directions (12-directions scheme). We denote the original ADC maps as *ADC*_20_ and those generated from these subsets as *ADC*_4_ and *ADC*_12_.

**Figure 2.**
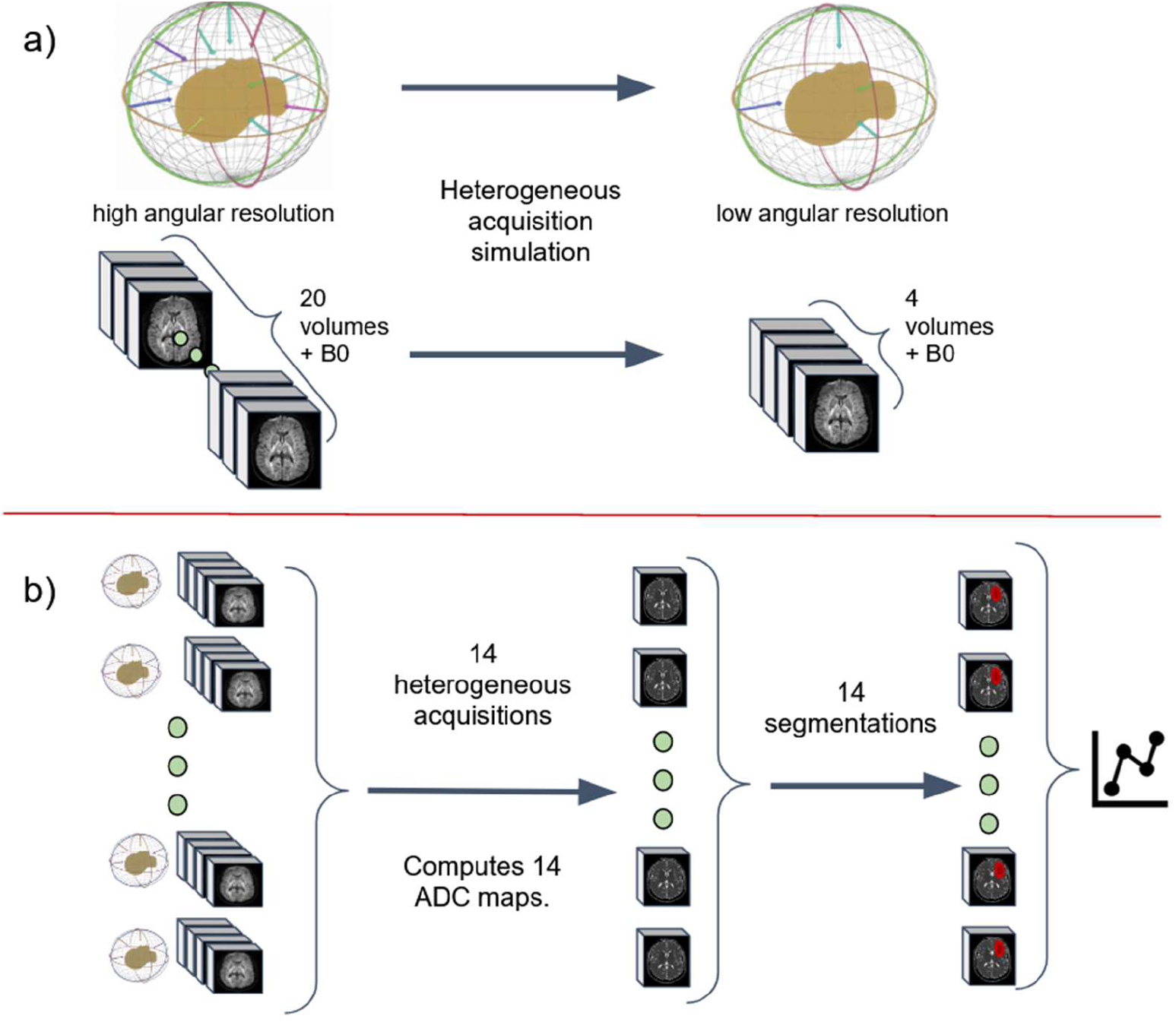
Heterogeneous protocol data generation procedure. (a) The subsampling procedure simulates a different acquisition protocol with less diffusion encoding directions. Each subsample consists in a new set of three orthogonal directions plus one random direction. Each new set can be considered as a new repetition where local orientation with respect to the subject changed due to head tilting or protocol difference. (b) 14 such heterogeneous simulated sets of diffusion encodings are generated, yielding 14 variations of ADC maps. The infarct core mask is computed using a fixed 620 × 10^−9^*m*^2^/*s* threshold. Spurious regions are removed using morphological operations.

In addition to the *ADC*_4_ and *ADC*_12_ maps, our study also included the generation of several *ADC*_4_ maps through an adjusted subsampling method. For each iteration, a new subset of diffusion gradient directions was chosen from the original high-angular resolution protocol, with the aim of selecting directions that were nearly equispaced. This method was designed to simulate different diffusion sampling schemes and evaluate their effect on ADC maps. By creating these varied *ADC*_4_ maps, we examined the influence of distinct gradient direction configurations on the ADC values. The process is illustrated in Figure 2, panel B, offering a visual representation of how different gradient direction choices affect ADC maps and, by extension, the assessment of stroke lesions. This analysis provided insights into how the selection of diffusion gradients might impact the precision and reliability of stroke lesion segmentations.

### Threshold-based lesion segmentation

In order to determine the presence of lesions, a segmentation process was carried out on the ADC maps, focusing only on the white matter (WM). To do so, the WM was first segmented using SynthSeg (Billot et al., 2023) on the T2-weighted images, and then registered to the DWI image space using the Elastix software (Klein et al., 2010). To generate the infarct core masks, the widely used DEFUSE criterion (Purushotham et al., 2015) was applied to all the volumes. This criterion is based on a single threshold at 620×10^−6^ mm^2^/s to delineate the stroke ischemic core. The study exclusively examined the WM, as this threshold has been reported to include numerous false positive lesion voxels in deep cortical GM and to overestimate the infarct core (Purushotham et al., 2015). Consequently, it is deemed unspecific and does not easily generalize to various datasets, as reported by (Khoury et al., 2019). To address these limitations, the resulting masks were further processed to reduce the number of incorrectly segmented lesions. This was achieved by applying a morphological opening operation, followed by a closing operation, using a 2×2×2 voxel-sized squared structuring element. The binary mask was then decomposed into 3D connected components using a 27-voxel isotropic neighbourhood to segment the spurious lesions. Finally, any small component with a volume lower than 1 mm^3^ was filtered out. The source code to replicate this DEFUSE thresholding procedure is available at **HTTPS://GITLAB.COM/TRANSLATIONALML/ASAP/DWISTROKE**.

## Statistical analysis

The aim of the primary analysis was to test whether ischemic core volume and location, as determined by the DEFUSE criterion, depended on DWI acquisition protocol. The aim of the secondary analysis was to test whether such acquisition protocol changes also modified associations between ischemic core volume and clinical scores.

### White Matter ADC Distribution

In this analysis, we compared the ADC distributions in white matter derived from the high-angular resolution *ADC*_20_ and the reduced protocol *ADC*_4_. We aimed to detect any consistent shifts in mean ADC values or alterations in the distribution’s spread, which could potentially impact the application of the DEFUSE-based segmentation criteria, particularly considering its dependence on specific ADC thresholds. To assess the differences between these two protocols, we employed the Wilcoxon signed-rank test, a non-parametric statistical test suitable for comparing paired samples. This test was chosen to evaluate the significance of differences in mean ADC values, as well as the variability (standard deviation), kurtosis, and skewness of the ADC distributions.

### Ischemic core volume

A Bland-Altmann analysis of the infarct volume was used to assess the non-parametric 95% range of agreement between the 20-directions and 4-directions acquisition schemes, that is, the interval within which 95 per cent of the differences in volume due to acquisition scheme are expected to fall.

We then used multivariate regression in order to evaluate the impact of acquisition protocol on infarct core volume while accounting for covariates. Due to non-Gaussianity, a beta mixed model (Brooks et al., 2017; Ferrari & Cribari-Neto, 2004) was fitted with (rescaled) infarct core volume as the response and a probit link function, the acquisition scheme (20-directions versus 4 directions, using only the first repetition) as the fixed effect of interest, Age and Sex as covariates, and a random intercept random effect per subject. Significance of fixed effects was assessed by a Wald test and profiled confidence intervals, and the significance of the acquisition scheme was confirmed by comparing differences in corrected Akaike Information Criterion (AICc) (Sugiura, 1978) between a model comprising acquisition scheme as a fixed effect and a model without this predictor, to evaluate their relative support (Burnham & Anderson, 2004) (R package bbmle). Finally, we evaluated both full and reduced model with mean absolute error (MAE) and relative mean absolute error (RMAE).

### Ischemic core location

For each patient, the infarct core location consistency was assessed by computing Jaccard coefficients between the ischemic core voxels delineated by the original acquisition protocol and each of the 14 sampled schemes, resulting in 323 × 14 = 4522 Jaccard values. Summary statistics were obtained for the study samples by computing the median, maximum, minimum, and interquartile range over all subjects. Computations were performed using Python 3.9 and the Scipy 1.9.2 package (Virtanen et al., 2020).

Clinical associations

We examined the correlation between ischemic core volume and three clinical variables obtained from the registry: NIHSS at admission, NIHSS at 24 hours, and 3-months modified Rankin Scale (mRS); this was performed separately for the ischemic core volume obtained from the twenty-directions diffusion scheme and the four-directions diffusion scheme. Because neither clinical variables nor ischemic core volume were Gaussian, as described above, we used Spearman rank correlations. In the case of missing clinical variables, we used only complete cases.

We confirmed this univariate analysis by using a multivariate Generalized Additive Model (GAM), with a beta response for the clinical variable (after rescaling to the (0,1) interval, a spline smoother on ischemic core volume(Wood et al., 2016), and correction for sex and age (R package mgcv). We compared the model fitted on 4-directions data to the model fitted on 20-directions data using 500 bootstrap resampling, each time computing the difference in AICc.

## Results

### White Matter ADC Distribution

In comparing white matter apparent diffusion coefficient (ADC) characteristics between the 4-directions and 20-directions diffusion schemes, we found clear differences in mean ADC values, variability, and distribution shapes.

The mean ADC for the 20-directions scheme was 864.97 μm^2^/s, lower than the 4-directions scheme, which had a mean of 872.94 μm^2^/s. The standard deviation of ADC values in the 20-directions scheme was 261.90 μm^2^/s, indicating less variation compared to 283.08 μm^2^/s in the 4-directions scheme. Kurtosis was higher in the 20-directions scheme with a mean of 24.22, compared to 18.48 in the 4-directions scheme, pointing to a more peaked distribution of ADC values. Skewness also increased in the 20-directions scheme, with an average of 3.85, versus 3.14 in the 4-directions scheme, suggesting a more skewed distribution.

The Wilcoxon signed-rank test showed significant differences in the standard deviation, kurtosis, and skewness of ADC between the schemes. The test for the standard deviation of ADC values resulted in a p-value < 7.15 ×10^−16^, for kurtosis, a p-value < 6.72 ×10^−16^ and for skewness, a p-value < 4.55×10^−16^. These results confirm significant differences between the diffusion schemes in terms of ADC variability, the peakedness, and asymmetry of its distribution. The data strongly suggests that the choice of diffusion scheme affects ADC measurement characteristics in white matter, with the 20-directions scheme showing lower variability but higher kurtosis and skewness compared to the 4-directions scheme.

### Ischemic core volume

WM volume averaged 480.22 ml (sd 54.36 ml, max 670.45 ml). For the infarct volume, in the 12-directions scheme, it averaged 6.07 ml (sd 11.32 ml, max 119.86 ml), compared to 3.97 ml (sd 10.85 ml, max 118.10 ml) across patients in the 20-directions scheme, and 7.88 ml (sd 11.05 ml, max 117.54 ml) for those in the 4-directions scheme.

Infarct volumes were overall over-estimated in the 4 and 12-directions schemes compared to the 20-directions scheme. Figure 3 panel a) shows for the 4-directions scheme, the average bias was 3.91 ml, with a range of differences from -10.88 ml to 26.47 ml. In comparison, for the 12-directions scheme, the average bias was 2.10 ml, with a range of differences from -2.17 ml to 23.03 ml.

**Figure 3.**
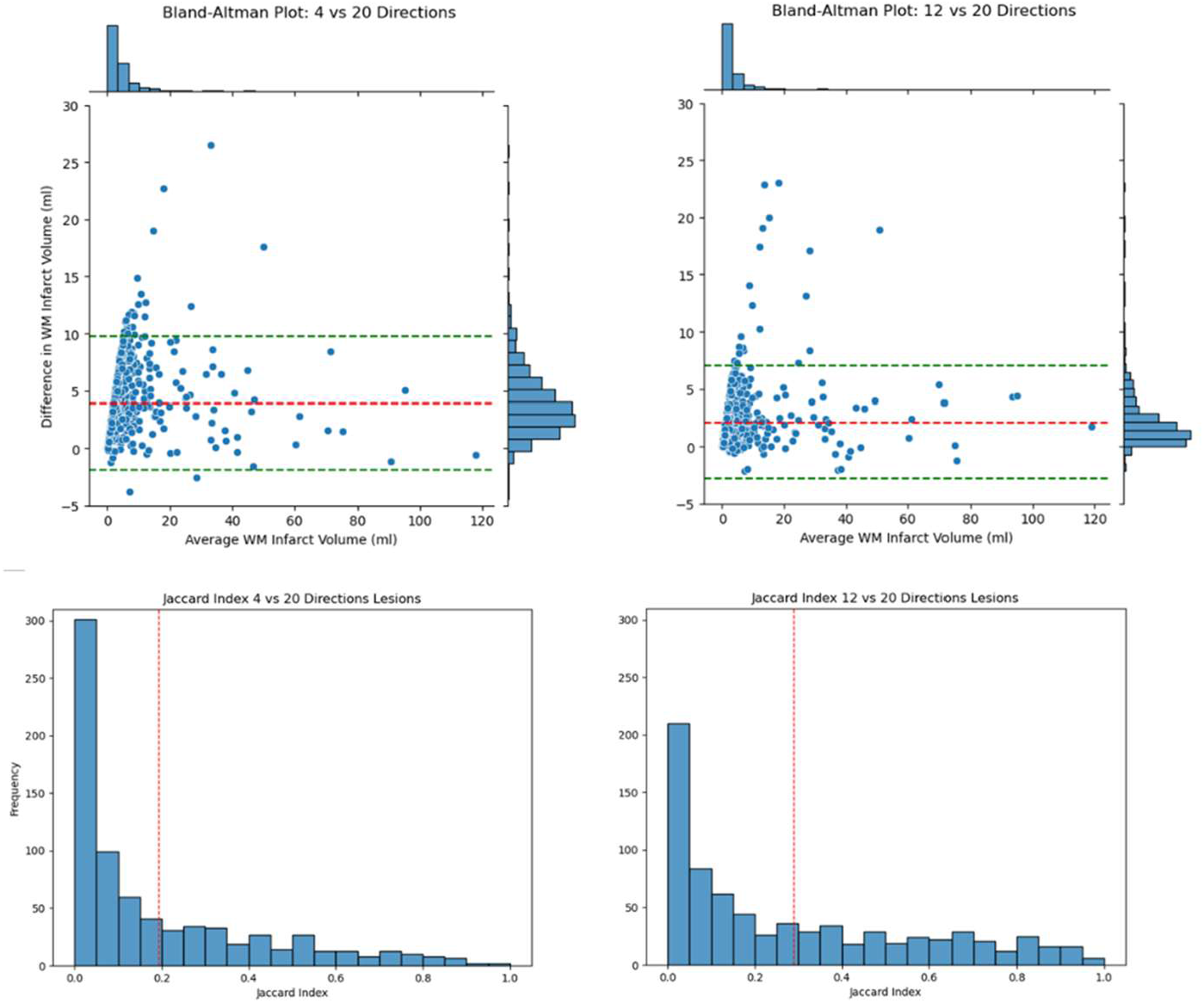
Top, Bland-Altman plot for the difference in infarct core volume between the 4-directions and the 20-directionsDWI protocol. The bias is 7.3 ml and the non-parametric 95% range of agreement is 0.4-23.6 ml. Bottom, Distributionof Jaccard index values across patients, indicating the degree of agreement on infart core voxel locations between the20-directions, and the 4-directions (left) and 12 directions (right) DWI protocols. Values of 0 indicates no voxels overlap between protocols, while 1 indicates all voxels overlap.

Multivariate regression on ischemic core volume confirmed the significant effect of the diffusion acquisition scheme. Specifically, using the 4-direction and 20-directions schemes, it results in a reduction of estimated ischemic core volume by an average of 0.396 ml (95% CI: −0.418 to -0.375, *z* = −36.0, *p* < 2 × 10^−1^). Compared to models excluding this scheme, the inclusion improves the AICc by 1012 points. In terms of accuracy, the full model with the acquisition scheme reports a Mean Absolute Error (MAE) of 2.9 ml and a Relative Mean Absolute Error (RMAE) of 0.22, improvements from 4.5 ml and 0.34, respectively, in models without the scheme.

### Ischemic core location

Figure 4 shows, for three random patients, the spatial overlap between infarct core locations computed using different directions. Complementary, Figure 4 quantifies the spatial overlap using the Jaccard coefficient for all subjects. The median (IQR) Jaccard index between the 20-directions acquisition and all 14 variations of the 4-directions scheme, across all subjects, was 0.22 (0.13-0.39)

**Figure 4.**
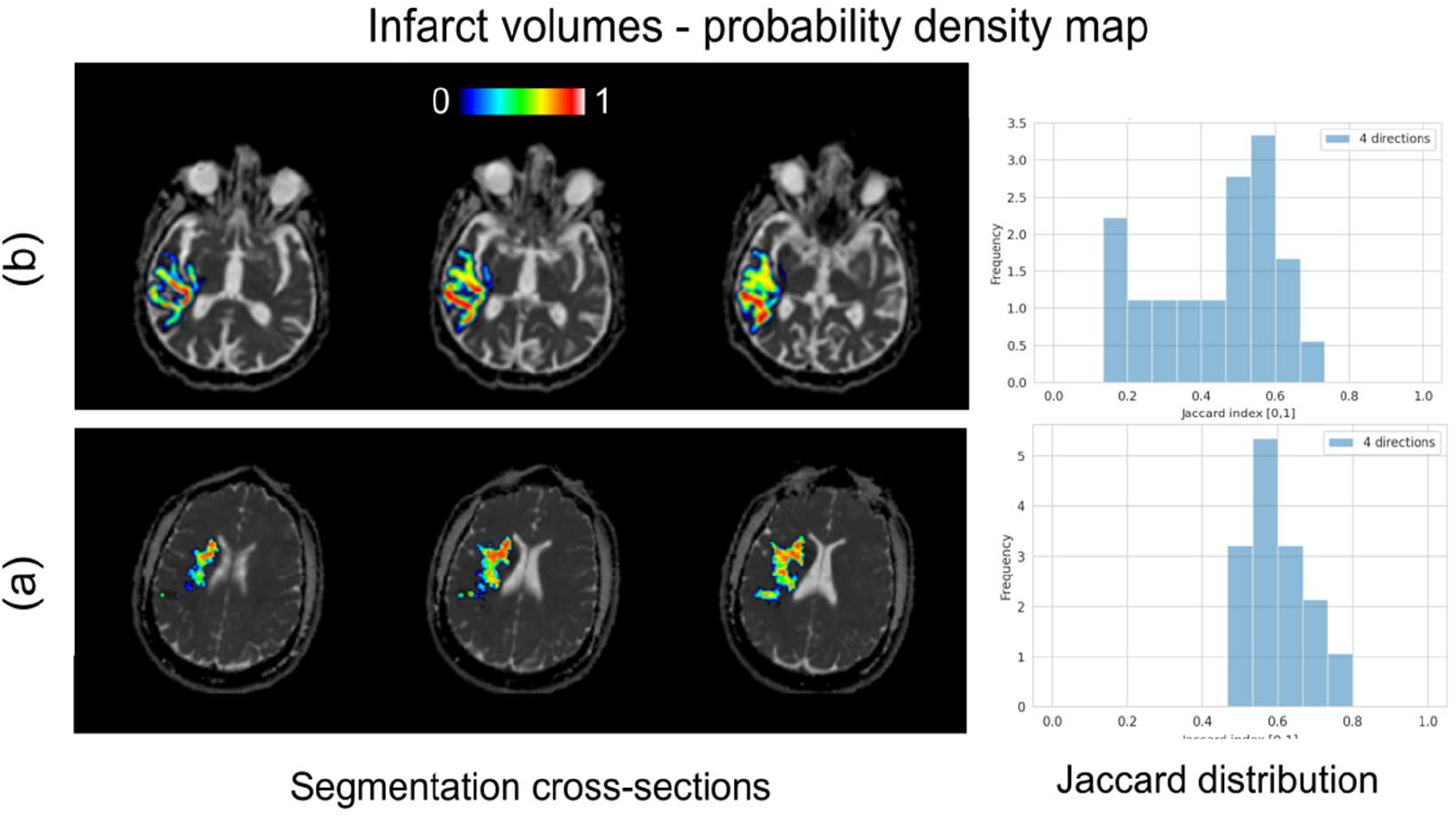
Infarct probability mask computed from the full set of fourteen subsamples for two randomly selected subjects with an infarct volume greater than 20 ml and a single apparent lesion in the white matter. For each subsample, the binary infarct mask was computed; all such infarct masks were added together, and then divided by the total number of subsampled schemes (14). Warmer colours indicate more consensus for infarct core voxels across the subsampled schemes, while voxels in cooler colours indicate that fewer subsampling schemes would assign them to the infarct core. Considerable variability in location can be observed across subsampled schemes and subjects as shown in the Jaccard distribution of the 14 repetitions on the right panel.

### Clinical associations

For NIHSS at admission (N=726), the demographics-only model had a significant effect of age (z=2.7, p=0.007) but not sex (z=1.5, p=0.14), and a low explained deviance (4.3%). Univariate Spearman correlation between ischemic core volume and NIHSS at admission was 0.257 (p=2.018×10^−12^, 95% CI 0.187-0.324) for 4-directions, and 0.471 (p<2.2×10^^-16^, 95% CI 0.413-0.526) for 20-directions imaging.

The multivariate regression model confirmed the significant effect of age and ischemic core volume for both 4-directions and 20-directions data, but the deviance explained was higher in the 20-directions model (29.3% vs 20.6%), and bootstrap analysis showed that the 20 directions model had much higher strength of evidence (0/500 resamples had higher deviance explained for the 4-directions than for the 20-directions data, and the median [IQR] delta-AICc for the 4-directions model was 37.7 [30.62,45.36]).

For NIHSS at 24 hours (N=726), the demographics-only model had a significant effect of age (z=2.7, p=0.008) but not sex (z=0.31, p=0.76), and a low explained deviance (6.0%). Univariate Spearman correlation between ischemic core volume and NIHSS at 24 hours was 0.227 (p=8.18×10^−10^, 95% CI 0.156-0.29) for 4-directions, and 0.436 (p<2.2×10^−16^, 95% CI 0.37-0.49) for 20-directions imaging. The multivariate regression model confirmed the significant effect of age and ischemic core volume for both 4-directions and 20-directions data, but the deviance explained was higher in the 20-directions model (24.6% vs 13.5%), and bootstrap analysis showed that the 20 directions model had much higher strength of evidence (0/500 resamples had higher deviance explained for 4-directions data, and the median [IQR] delta-AICc for the 4-directions model was 31.0 [23.9, 41.6]).

Finally, for modified Rankin scale at 3 months (N=726), the demographics-only model had a significant effect of age (z=3.6, p=0.0003) but not sex (z=1.4, p=0.15), and a low explained deviance (11%). The Univariate Spearman correlation between ischemic core volume was 0.06 (not significant at p=0.090, 95% CI 0.00-0.140) for 4-directions, and 0.27 (p=2.8×10^−13^, 95% CI 0.20-0.34) for 20-directions imaging. The multivariate regression model confirmed the significant effect of age and ischemic core volume for both 4-directions and 20-directions data, and the deviance explained was similar in the two models (20.7% in 20-directions, 16.2% in 4-directions); bootstrap analysis showed that the 20-direction models had slightly higher strength of evidence (29/500 resamples had higher deviance explained in the 4-directions model, and the median [IQR] delta-AICc for the 4-directions model was 10.5 [6.4,15.4]).

## Discussion

ADC maps derived from DWI data are used to delineate the infarct core in acute stroke. Using 726 retrospective patients, our aim was to investigate the effect of diffusion protocol changes on the estimation of infarct core volume, location, and association with clinical scores. When using a threshold-based approach, as commonly done in clinical trials, we showed that a 4-directions scheme overestimated infarct volumes compared to a 20-directions scheme (average 7.8 ml, range -1 ml to + 33 ml), a difference which held in multivariate analysis when correcting for age and sex. Finally, we showed that correlations between infarct volume and clinical scores were weaker in the 4-directions scheme than in the 20-directions schemes, and that such differences persisted in multivariate analysis correcting for sex and age: Spearman correlation in acute NIHSS was lowered to 0.25 from 0.47, 24 hours NIHSS was lowered to 0.22 from 0.43, and correlation with 3-months was lowered to 0.06 (not significant) from 0.27.

Our study demonstrates that differences in diffusion encoding, stemming from protocol changes (which may also be related to head tilting), significantly affect the ADC distribution in white matter, particularly the distribution’s spread. We corroborated athat the infarct location changed importantly between the different diffusion acquisition protocols, with a median Jaccard index of 0.22 across patients and diffusion schemes showing poor agreement with the 4-directions scheme yielding a heavier left tail for the ADC distribution. Such changes have considerable implications on the determination of infarct core volume and location when using thresholding methods such as those in DEFUSE. These alterations can introduce bias in multi-centric trials, as infarct sizes and locations may be skewed due to diffusion protocol differences or patients’ relative head positions. A key observation is the leftward shift in the ADC distribution for the 20-directions scheme, indicating a concentration of energy towards lower ADC values. This shift could elucidate or contribute to the heightened segmented threshold, which is pivotal given that the threshold for segmentation is around 620 *μm*^2^/*s*. The increased skewness, in particular, implies a longer tail of lower ADC values, whereas the elevated kurtosis points to a sharper peak closer to this lower threshold. These shifts in the ADC characteristics could have implications for the precision of white matter lesion segmentation, potentially leading to differences in the estimated extent of lesioned tissue when employing different diffusion schemes.

While our findings suggest that employing a higher number of directions in DWI acquisitions can offer more accurate measurements of infarct volumes, as these better reflect clinical impact, this may result in longer acquisition times and impair clinical adoption; however modern sequences and fast acquisition/reconstruction schemes can mitigate this increase, bringing acquisition times down to an acceptable range (around 4 minutes for the 20-directions EPI protocol used in this study, which could be further sped up).

Our study, however, has several limitations. First, while we focus on differences in diffusion directions, other protocol changes such as slice thickness specification, flip angle, etc. would also likely contribute to differences in volume, location, and clinical associations. Our results therefore represent a conservative estimate of protocol effects in stroke workups. Second, our low-angular resolution protocol is simulated from patient data rather than acquired anew on the same patients. While this is due to ethical constraints in acute imaging, it is possible that our low-angular data does not faithfully reflect actual clinical protocols. Finally, we excluded many patients due to either missing data or image quality, principally motion artefacts. This is a potential source of bias, and it is therefore possible that our findings are specific to patients that lays relatively still in the scanner and would fail to generalize to patients that tend to move more – though it is difficult to speculate as to the differential effect of motion on our results.

In conclusion, our study showed that DWI acquisition protocol differences can substantially alter apparent infarct volume, infarct location, and clinical associations, and emphasizes the need for careful specification of diffusion acquisition protocol to improve the accuracy and consistency of acute stroke workups, ultimately enhancing patient care and outcomes.

## Abbreviations

DWI: Diffusion-weighted magnetic resonance imaging
PWI: Perfusion-weighted imaging
ADC: Apparent diffusion coefficient
NIHSS: National Institutes of Health Stroke Scale/Score
mRS: Modified Rankin Scale.
EPI: Echo-planar imaging
TE: Echo time
TR: Repetition time
IQR: Interquartile Range
AICc: Corrected Akaike Information Criterion

## Acknowledgments

We thank interventional radiologists Dr Bruno Bartolini and Dr Steven Hajdu for helpful discussions of this work.

## Notes

### Competing Interest Statement

The authors have declared no competing interest.

